# PBRM1 is a vulnerability in ARID1A deficient multicellular tumor spheroids

**DOI:** 10.1101/2022.02.21.481305

**Authors:** Tomali Chakravarty, Kathleen Diep Tran, Dinesh Dhamecha, Tyrus Perdue, Jada L. Garzon, Jyothi U. Menon, Arnob Dutta

## Abstract

ARID1A, a subunit of SWI/SNF, has been shown to play a major role in recruitment of the chromatin remodeler to enhancers for transcriptional regulation. Mutations inARID1A have been found in various cancers, many of which form solid tumors. Recent studies have revealed vulnerabilities in cells lacking *ARID1A*, specifically ARID1B, an ortholog and mutually exclusive subunit, in 2D cell culture. However, identification of vulnerabilities within SWI/SNF for loss of ARID1A in a multicellular tumor spheroid, that mimic in vivo condition within tumors, has not been explored. Here we show in the absence of ARID1A in a MTS model, ARID1B continues to be a vulnerability but we have identified PBRM1 as a new vulnerability within SWI/SNF. Levels of ARID1B and PBRM1 are elevated on loss of ARID1A. Further, reduction of ARID1B and PBRM1 protein levels, decreases cell survival and reduces induction of several hypoxia regulated genes in ARID1A deficient MTSs. Our studies have identified PBRM1 as a new vulnerability in ARID1a deficient cancers and which provides a new target for therapeutic strategies.

## Introduction

The SWI/SNF complex is a multi-subunit, ATP-dependent chromatin remodeler that functions in gene regulation and modulation of chromatin architecture^1,2^. SWI/SNF is evolutionarily conserved in eukaryotes and in humans, there exist three distinct assemblies: BRG1/BRM-associated factor complexes (BAFs), polybromo-associated BAF complexes (PBAFs), and non-canonical BAFs (ncBAFs).^3,4^ All assemblies contain an ATPase subunit, either BRG1 or BRM, and several core subunits, including BAF60a, BAF155/BAF170, INI1 and BAF57 (INI1 and BAF57 found exclusively in BAF/PBAF complexes). Each assembly can be recognized by exclusive subunits: BAF contains ARID1A or ARID1B, PBAF incorporates ARID2, BRD7 and PBRM1 and ncBAF consists of GLTSCR1/L and BRD9. The heterogeneity of SWI/SNF allows for its selective genomic targeting at enhancers and/or promoters by each complex^5–10^. Therefore, SWI/SNF complexes are involved in regulating many processes, including development^11–14^, differentiation^11,12^, and DNA repair.^15,16^

SWI/SNF complexes have emerged as a major focus of attention because of the mutational frequencies in several specific subunits that have been found in a range of human diseases. Alterations and inactivation of genes encoding subunits of SWI/SNF have been implicated in over 20% of cancers^17,18^, linked to neurological disease^19^, and associated with intellectual disability and autism spectrum disorders.^20,21^ Of the subunits of SWI/SNF, it is not only the catalytic subunit (ATPase), BRG1 or BRM, that are highly mutated, but also include mutations in complex specific subunits like ARID1A, a subunit of BAF which has a role in targeting the complex to tissue-specific enhancers and maintaining chromatin accessibility at these locations^22^. ARID1A mutations are found in over 50% of ovarian clear cell carcinoma and approximately 30% of ovarian endometroid carcinomas ^23,24^, while in colon cancers, ARID1A mutations are present in about 10% and caused by mismatch defects.^25^ These mutations of ARID1A have indicated that ARID1A is a tumor suppressor as in many cancers, loss of this protein has been shown to be a driver of tumor formation due to dysregulation of gene expression.

There are two homologous subunits of ARID1A, ARID1B and ARID2, which are used to assemble the SWI/SNF complexes; ARID1A or ARID1B members of the BAF complex and ARID2 which is a part of PBAF complexes. ARID1A and ARID1B are orthologues that share 60% identity in protein sequence and are mutually exclusive members of the BAF complex. Although ARID1A and ARID1B are co-expressed, ARID1A is more abundant than ARID1B which could account for the higher rate of mutations found in ARID1A than ARID1B in cancers^6^. Interestingly, the functions of ARID1A are also observed for ARID1B, but only in the absence of ARID1A.^6^ Studies have shown in both ovarian and colorectal carcinomas that are deficient in ARID1A that loss of ARID1B decrease cell viability^26,27^, but there is a lack of small molecule inhibitors that alter ARID1B function, making ARID1B a difficult target. Our study has extended to the other complexes that have not been well studied in cells lacking ARID1A.

As these studies were performed in two-dimensional (2D) monolayers, we asked if these results could be recapitulated in three-dimensional (3D) cell culture as ARID1A has a role in tumor formation. 3D cell culture behaves and reflects more accurately of the microenvironment where cells reside.^28,29^ We decided to use an innovative 3D model to study the vulnerabilities of cells deficient in ARID1A. We employed the colorectal carcinoma cell line, HCT116 (expresses protein ARID1A) and HCT116 ARID1A-/- (no expression of ARID1A protein), as other studies have looked at vulnerabilities in this cell line, express all forms of SWI/SNF and are able to form spheroids.^30,31^ The composition characterization and activity of unique subunits of SWI/SNF have not been explored greatly in 3D models. In this study, we characterized the three complexes of SWI/SNF in 2D and 3D models and revealed new vulnerabilities in multicellular tumor spheroids (MTSs) lacking ARID1A. We show that ARID1B-BAF and PBAF were present at elevated levels in the spheroids lacking ARID1A and loss of ARID1B and PBRM1 decreased viability of ARID1A-deficient MTSs. ARID1B and PBRM1 were also important for induction of genes on loss of ARID1A. Our studies suggest that PBRM1 is a new therapeutic target for tumors with ARID1A mutations.

## Results

### SWI/SNF composition is altered in cells lacking ARID1A

We began by characterizing SWI/SNF in 2D monolayer cultures in two colorectal carcinoma lines, HCT116 (wild-type) that expresses ARID1A, and in the same cell line, HCT116 ARID1A/- ((Horizon Discovery) generated by a knock-in of a premature stop codon (Q456*) on both chromosomes and expresses no ARID1A protein) that has lost expression of homozygous ARID1A. We examined nuclear protein levels of core subunits, the catalytic subunit BRG1 and several unique subunits of each assembly (Figure 1A). All three assemblies were present in HCT116. In HCT116 ARID1A-/-, protein levels of core subunits were not significantly altered compared to HCT116. ARID1A protein was not present as expected inHCT116 ARID1A-/- and we observed an increase in ARID1B protein levels in the mutant compared to the wild-type. Interestingly, we also observed an increase in expression levels of PBAF member PBRM1 and GLTSCR1, subunit specific to the ncBAF form of SWI/SNF (Figure 1A), suggesting that these two assemblies may play important roles in cell lines lacking *ARID1A*. To further validate our findings, we performed immunoprecipitation to purify endogenous SWI/SNF complexes through the catalytic subunit, BRG1, followed by mass spectrometric analysis. This analysis revealed that levels of association of core subunits with BRG1 was slightly decreased on loss of ARID1A (Figure 1B). In HCT116 ARID1A-/- cells, the BAF complex predominantly was represented by presence of ARID1B. Interestingly, increased association of PBAF complex specific subunits PBRM1 and BRD7, as well as ncBAF specific subunits members GLTSCR1 and BRD9, with BRG1 was observed. The elevated levels of PBAF and ncBAF in cells lacking *ARID1A*, may suggest that cells may display greater dependence on these complexes in in addition to ARID1B for gene expression and survival.

**Figure 1:**
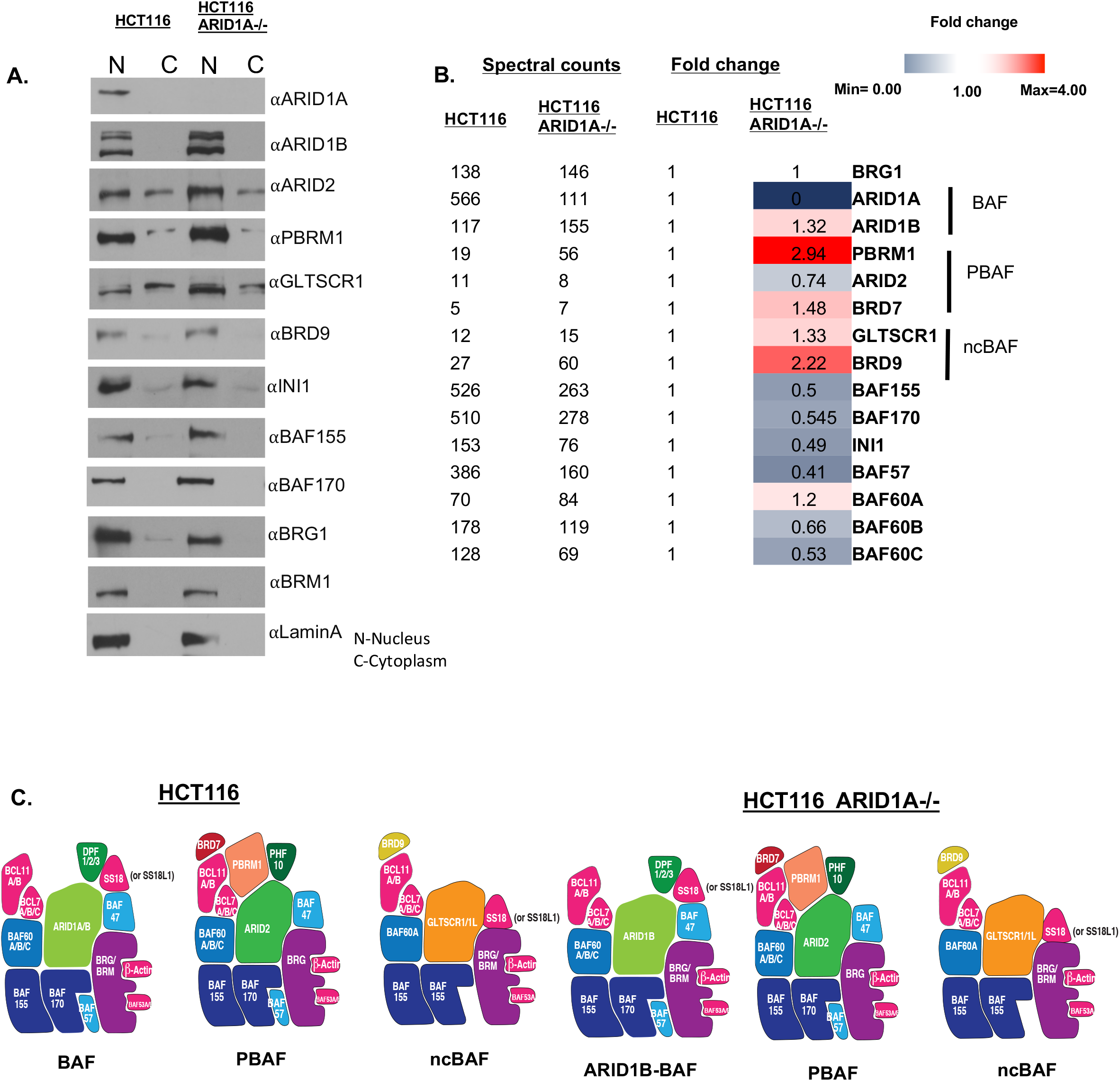
Balance of SWI/SNF is altered in HCT116 ARID1A-/-cells. **A)** Western blot images showing SWI/SNF protein levels in nucleus and cytoplasmic fractions from HCT116 and HCT116 ARID1A-/- cells grown in monolayer. **B)** Mass spec data from HCT116 and HCT116 ARID1-/- cells showing altered balance of SWI/SNF complex. In the table are the total spectral counts of different proteins from BRG1 immunoprecipitation in both cell lines. The fold changes in protein levels were calculated by normalizing the levels of protein to HCT116 levels set to 1. **C)** Human SWI/SNF exists in three distinct assemblies-BAF, PBAF and nc BAF. HCT116 cells contain all the three complexes. In HCT116 ARID1A-/-, absence of ARID1A causes BAF complex to be solely dependent on ARID1B.

### Loss of ARID1B and PBRM1 decreases survival of multicellular tumor spheroids lacking ARID1A

While previous studies have shown that cells lacking ARID1A are dependent on its mutually exclusive ortholog ARID1B, the roles of other SWI/SNF complexes in cells lacking ARID1A, have not been evaluated. Since we found that significant increases in PBAF member PBRM1 levels and ncBAF member BRD9, we next determined the roles of these subunits in survival of ARID1A^-/-^ cells. Small interference RNA (siRNA) was used to decrease levels of ARID1B, PBRM1, BRD9, BRG1, with a non-target siRNA used as control in both HCT116 and HCT116 ARID1A-/- cells. Reduction in protein levels on siRNA mediated knockdown was verified by western blotting (Supplementary Figure 1A). Viability of HCT116 non-target treated siRNA was used as control and set to 100%. Knockdown of subunits, representative of each form of SWI/SNF in HCT116 showed no changes in cell viability (Supplementary Figure 1B). Loss of BRG1 significantly reduced viability in both cell lines. In HCT116 ARID1A-/-, we observed as has been shown earlier, that loss of ARID1B decreased cell viability^26,27^ (Supplementary Figure 1B). Contrary to expectations, though the levels of PBRM1 and BRD9 were elevated in cells lacking ARID1A, knockdown of these proteins did not significantly affect cell survival on loss of ARID1A. Loss of ARID1A is often associated with ovarian and endometrial carcinomas, both of which form solid tumors. Solid tumor environment significantly differs from those in cells cultured in a 2D culture system. We surmised that a possible reason for not observing expected effects on cell survival on loss of PBRM1 and BRD9 could be due to their contributions in the formation and maintenance of tumor environments on loss of ARID1A, which were lacking in a 2D culture system.

We turned our attention to a use of a well-defined multicellular tumor spheroid model which helps recreate an in vivo tumor microenvironment in which cells experience decreasing gradients of nutrients and oxygen from the outer surface to inner core of the spheroid.^32,33^ Also, the transcript profiles of MTSs, mirror more closely *in vivo* tumors than 2D cultures.^34,35^ To evaluate the effects of loss of ARID1B (BAF), PBRM1 and BRD7 (PBAF), BRD9 (ncBAF) and INI1 (found in BAF/PBAF) on growth and survival of MTSs lacking ARID1A, we first generated MTSs from HCT116 and HCT116 ARID1A-/- cells (Supplementa Figure 2) where these subunits were knocked down by siRNA prior to formation of spheroids. Non-target siRNA was used as controls. Loss of subunits knocked down by siRNA treatment was confirmed by western blotting (Supplementary Figure 5). In order to determine populations of live and dead cells within spheroids, MTSs were stained using calcein AM (green fluorescence), that mark metabolically viable cells and ethidium homodimer-1 staining (red fluorescence), that mark non-viable cells, imaged by confocal microscopy. Confocal imaging of HCT116 showed that cells were spherical in shape and displayed predominantly green staining indicating no major growth defects (Figure 2A). Both loss of PBRM1 and ARID1B mirrored WT spheroids with no observable defects in MTS formation or growth. Though loss of BRD7 and BRD9 formed spherical spheroids, some increased death as indicated by red staining in the core, was observed. Loss of INI1 however resulted in significant defects in MTS formation, with only lose aggregation of cells being observed and increased loss of viable cells as determined by increased red staining observed in spheroids (Figure 2A). This is most likely because, loss of INI1, a core subunit used by two of the three forms of SWI/SNF can be expected to significantly impact gene regulation and survival of cells, which in turn would affect the ability of cells to form MTSs. On evaluation of MTSs formed from cells lacking ARID1A, we did not observe significant changes in spheroid shape and viable cells as compared to control MTSs with ARID1A. However, loss of ARID1B, PBRM1 and BRD7 in MTSs formed from cells lacking ARID1A, displayed significant defects in spheroid formation, and associated decrease in viability as compared to MTS lacking ARID1A alone. Loss of BRD9 however did not impact formation or viability in MTSs without ARID1A.

**Figure 2:**
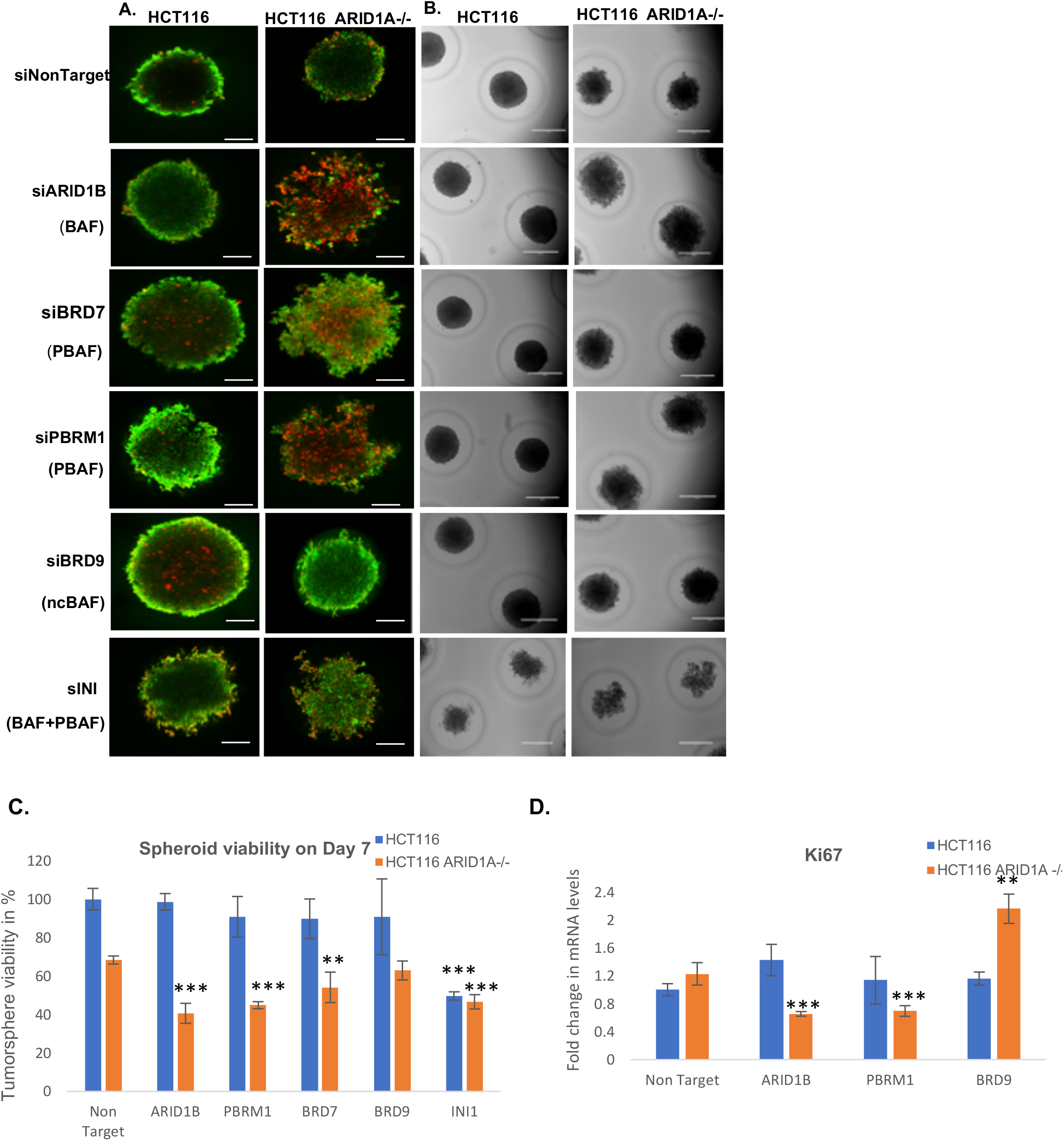
Knockdown of ARID1B and PBRM1 affects viability in Multicellular Tumorspheres lacking ARID1A-/-. **A)** Cell viability visualization of siRNA treated Tumorspheres, using a live/dead staining with ethidium homodimer-1 and calcein AM. Viable cells appear as green, while dead cells appear as red. HCT116 and HCT116 ARID1A-/- cells were treated with siRNA against subunits of SWI/SNF and allowed to form tumorspheres for 7 days. Images represent a section of Z stack taken using confocal microscope. Scale bars represent 100 μm. **B)** Brightfield images of tumorspheres treated with siRNA taken using EVOS microscope. Scale bars represent 400 μm.. **C)** Quantification of Spheroid Viability with Promega 3D Cell Titre GLO. Error bars represent SD of three independent experiments. **D)** qRT-PCR analysis of the mRNA levels of KI67 in HCT116 and HCT116 ARID1A-/- MTS. Error bars represent SD of three independent experiments. ***<0.05

To quantitatively assess the viability of spheroids where unique subunits of SWI/SNF were depleted, we used an MTS assay optimized for spheroids (Promega). We examined viability of the spheroids on Day 3 and Day 7. Viability of HCT116 non-target treated siRNA was used as control and set to 100%. Loss of INI1 significantly reduced viability in MTSs both with and without ARID1A at both time points (Figure 2B). However, loss of any single subunit in MTSs with ARID1A, displayed no significant changes in viability. Loss of ARID1A alone showed a slightly decreased viability as compared to MTSs with ARID1A. Interestingly, further loss of ARID1B at Day 3, in spheroids without ARID1A, displayed significant loss of viability, to levels similar to those observed for cells lacking INI1(Supplementary Figure 3B). However, at Day 7, loss of ARID1B did not show a difference to Day 3 but loss of PBRM1 in spheroids without ARID1A slowly decreased cell viability from Day 3 to Day 7(Figure 2C). Loss of BRD9, between both time points, showed no defects in viability (Supplementary Figure 3B). In agreement with decreased viability in ARID1A^-/-^ MTSs lacking ARID1B and PBRM1, levels of proliferation marker Ki67 was also reduced (Figure 2D). These results cumulatively show that ARID1B and PBRM1 are required for survival of ARID1A-deficient spheroids and loss of PBRM1 is also synthetic vulnerability in ARID1A cancers in addition to ARID1B.

### ARID1A^-/-^ HCT116 MTS rely heavily on ARID1B-BAF and PBAF

Our results show that loss of ARID1B and PBRM1 result in decreased viability in MTSs lacking ARID1A. Observations from purification of SWI/SNF from 2D cell culture shows elevated levels of ARID1B and PBRM1 when cells lose ARID1A. However, since cellular environments can vary significantly between cells found in 2D vs those within MTSs, we sought to define the composition of SWI/SNF within MTSs with/without ARID1A. We performed an immunoprecipitation using BRG1 antibody in HCT116 and HCT116 ARID1A^-/-^ spheroids followed by mass spectrometric analysis of the complexes. Spectral counts of SWI/SNF subunits from HCT116 normoxia samples were used as control and set to 1. All fold changes were calculated with respect to HCT116 spectral counts. (Figure 3A). We observed core subunits to continue to be consistent between both 2D and MTS in HCT116. Interestingly, we observed a significant decrease in ARID1B within MTSs compared to monolayer cells (Figure 3A) suggesting that MTSs may not depend on ARID1B form of BAF. The levels of members of other complexes were not significantly altered. When we examined HCT116 ARID1A-/- spheroids, the decrease in protein levels of core subunits observed in 2D cells was not recapitulated within MTSs (Figure 3A). Levels of ARID1B-containing BAF was slightly increased in ARID1A^-/-^ MTSs compared to MTSs with ARID1A. This observation is in line with results showing an increase in ARID1B, on loss of ARID1A and importance of ARID1B in cell survival when ARID1A was lost^36,37^. Focusing on the other complexes, we found an increase in levels of PBAF members on loss of ARID1A with a significant increase in PBRM1 levels. ncBAF members were also moderately increased on loss of ARID1A with the exception of BRD9. To further confirm incorporation of subunits into their respective complexes we separated SWI/SNF complexes by glycerol gradient centrifugation followed by western blotting (Figure 3B). In HCT116, the three assemblies were well separated across the gradient and peaked in distinct fractions, ncBAF in fractions 11-14, BAF in fractions 14-18, and PBAF in fractions 17-20. BRG1 broadly occupied fractions but with the most concentrated in fractions associated with BAF, suggesting BAF may be the predominant complex present in HCT116 spheroids. In HCT116 ARID1A-/- MTSs, BRG1 was more equally divided among the fractions, suggesting the increased abundance and possible elevated roles of PBAF and ncBAF, along with ARID1B BAF, when ARID1A was lost. Mirroring results from our mass spectrometric analysis, we observed increased levels of ARID1B and PBAF members, PBRM1, ARID2, and BRD7. Interestingly, the level of BRD9 within ncBAF was reduced in ARID1A^-/-^ MTSs and shifted to being present in lower fractions (3-6), perhaps indicating free subunits that have not been incorporated within the complex (Figure 3B). This further confirms the low spectral counts observed from BRG1 IP inHCT116 ARID1A-/- MTSs, suggesting that either BRD9 may be not associating into SWI/SNF or has additional independent roles from those within ncBAF. Our results of increased levels of ARID1B and PBRM1 within MTSs lacking ARID1A further support our observations that these proteins are important for cell survival on loss of ARID1A.

**Figure 3:**
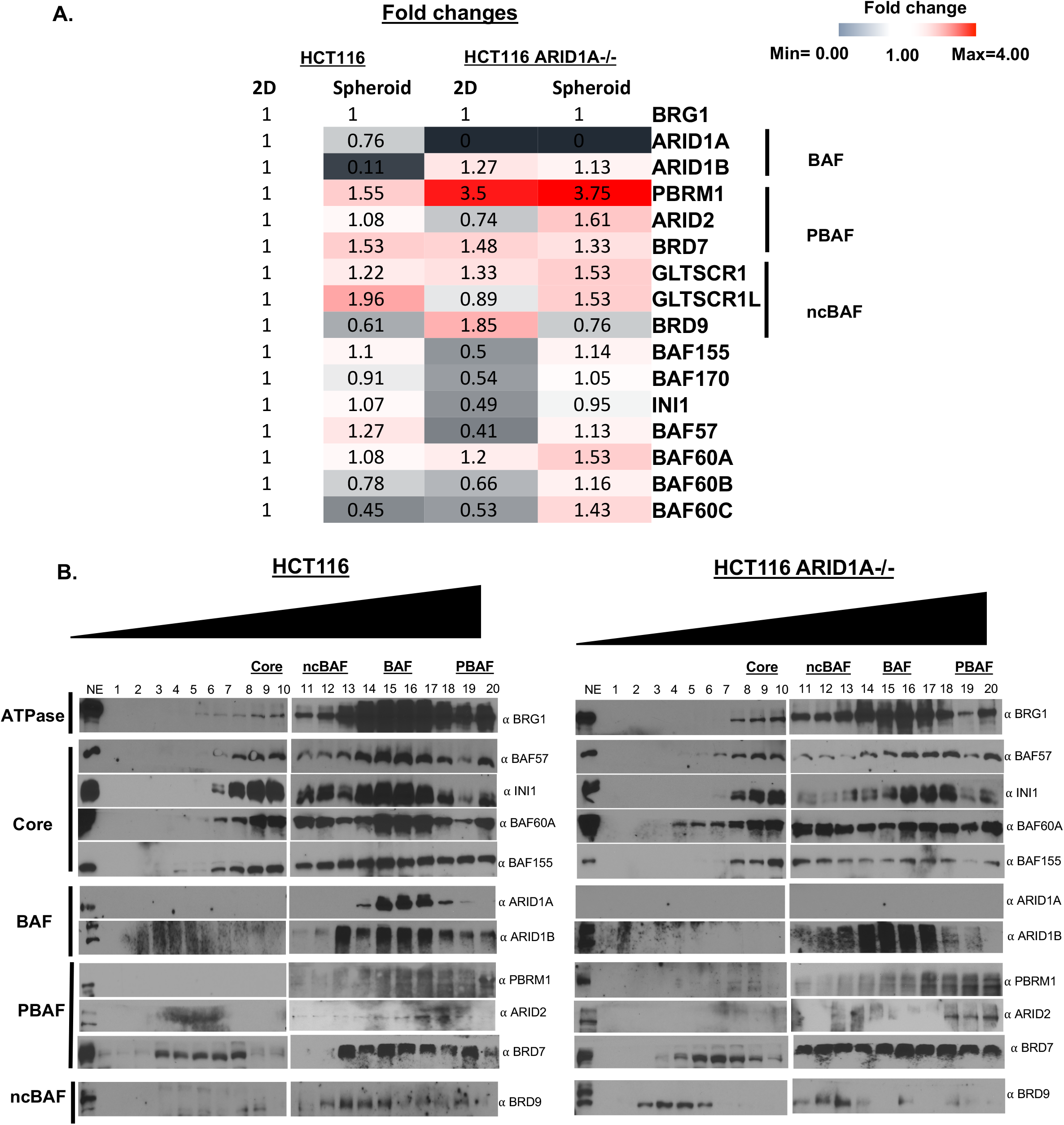
SWI/SNF composition changes in HCT116 ARID1A-/- Multicellular Tumorspheres. **A)** Mass spec data from HCT116 and HCT116 ARID1A-/- cells showing altered balance of SWI/SNF complex. Cells were grown as monolayer culture or allowed to form MTS for 72 hours. BRG1 immunoprecipitation was performed from both cell lines grown as monolayer culture or MTS. The fold changes in protein levels were calculated by normalizing the levels of protein to HCT116 levels set to 1. **B)** Western blots analysis of SWI/SNF complex subject to glycerol gradient. Nuclear proteins were isolated from 3-day old HCT116 and HCT116 ARID1A-/- spheroids. 1mg of nuclear protein was loaded on10 ml of 10-30% glycerol gradient column and spun at 35000 rpm for 18 hours. 500 μl fractions were collected, and the 20 fractions were subjected to SDS PAGE.

### Loss of PBRM1 and ARID1B affects gene expression within ARID1A^-/-^ MTSs

Multicellular tumor spheres consist of a milieu of cells experiencing varying oxygen saturation, depending on the location within the MTS, with peripheral cells located on the outside exposed to the most oxygen and a decreasing gradient of oxygen availability for cells in the interior of the MTS, resulting in a hypoxic core (Figure 4A). SWI/SNF has been shown to be an important player in the induction of hypoxic genes in 2D monolayers.^38–41^ Given the increased death we observed in the core of ARID1A^-/-^ MTSs lacking ARID1B and PBRM1 (Figure 2A), we surmised that these subunits may play important roles in hypoxic gene expression when ARID1A was lost. Spheroids formed from HCT116 and HCT116 ARID1A-/- cells treated with siRNAs against ARID1B, PBRM1, BRD9 and a non-target were analyzed for levels of hypoxia responsive genes, PAI1 and EPO that have been shown previously to be SWI/SNF dependent. Loss of ARID1B and PBRM1 reduced induction of PAI1 and levels of EPO in spheroids lacking *ARID1A* (Figure 4B). Levels of GLUT1, a HIF1 target gene that is known to be SWI/SNF independent, did not significant change, as compared to loss of ARID1A alone (SupplementaryFigure 4A). This suggests that the elevated levels of ARID1B and PBRM1 within MTSs lacking ARID1A, do indeed play important roles in gene expression and are important to compensate for loss of ARID1A.

**Figure 4:**
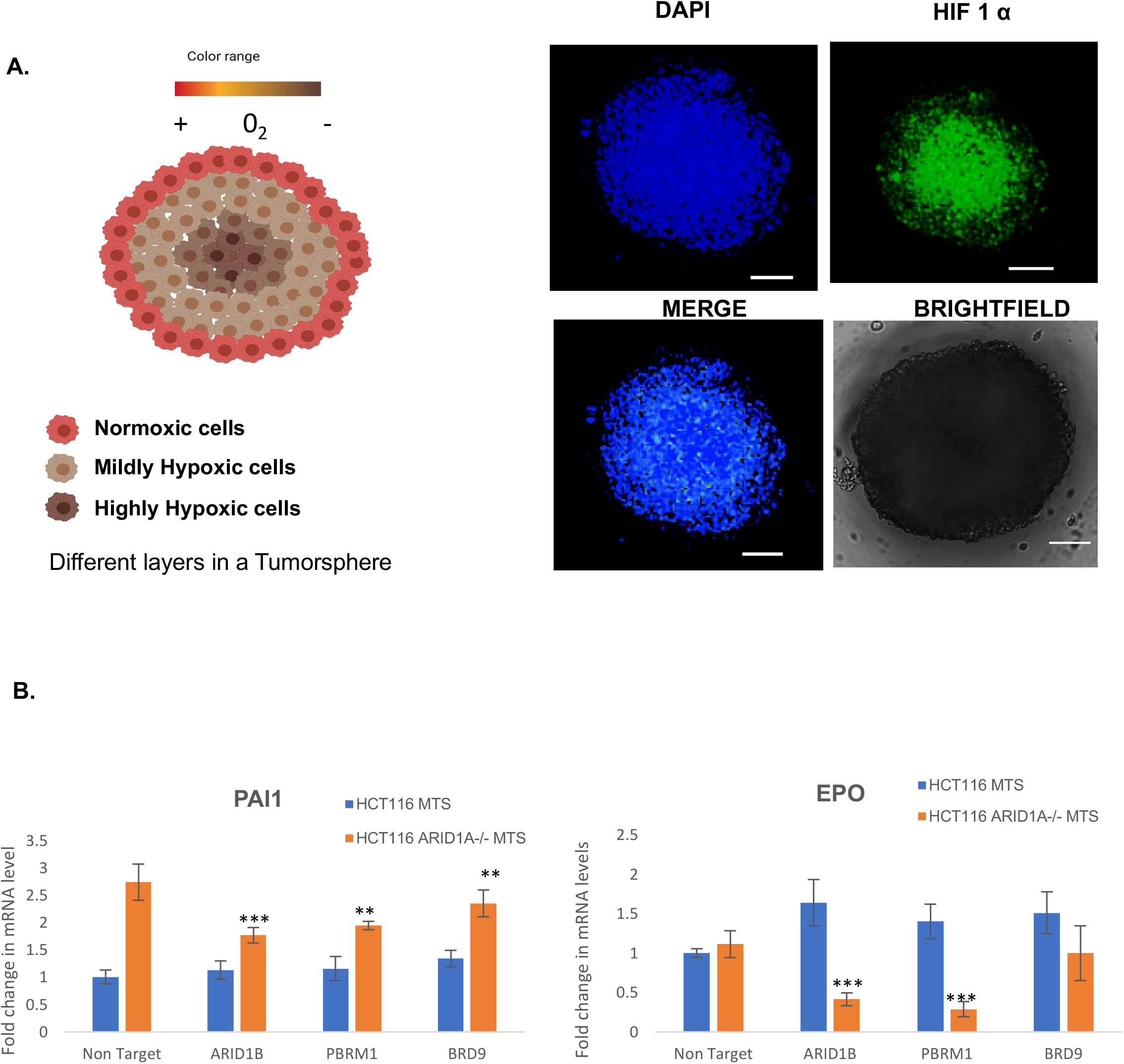
Knockdown of ARID1B and PBRM1 in HCT116 ARID1A-/- MTS affects hypoxic gene regulation. **A)** Immunofluorescence study for the hypoxia marker HIF 1α (green) shows hypoxia development within HCT116 spheroids. Spheroids were stained using primary antibody against HIF1α and Alexa Fluor 488 secondary antibody. DAPI was used to stain the nucleus. Images represent a section of the spheroid as captured by Zeiss confocal microscope. The Scale bar represents 100μm. **B)** qRT-PCR analysis of the mRNA levels of Hypoxia responsive genes PAI1 and EPO in HCT116 and HCT116 ARID1A-/- MTS. Cells were treated with siRNA against specific subunits and allowed to form MTS for 72 hours. RNA was isolated from the spheroids and mRNA quantified to calculate fold changes in induction. Error bars represent SD of three independent experiments. **<0.05

### ARID1A^-/-^ HCT116 MTS are sensitive to PFI-3

We have shown that loss of ARID1B and PBRM1 increase vulnerability of MTSs lacking ARID1A^-/-^. Earlier we had shown that loss of PBRM1 in ARID1A^-/-^ cells, resulte in increased death within MTSs formed from these cells. To confirm if inability of PBRM1 to bind to DNA affects cell viability within MTSs formed from cells lacking ARID1A, we used a PBRM1 inhibitor PFI-3.^42,43^ 3-day old HCT116 and HCT116 ARID1A-/- MTSs were grown in increasing concentration of PFI-3, for an additional 72 hours and visualized by confocal microscopy and analyzed for viability. Treatment of MTSs containing ARID1A, did not show significant death at low concentrations of the drug but were affected at high concentrations above 400 μM as evidenced by increase in red staining within the MTSs (Figure 5A). However, MTSs formed form cells lacking ARID1A, the red fluorescence could be detected at 75 μM, and increased at 150 μM (Figure 5A). At approximately 400 μM, the size of the spheroid has decreased significantly, and the red fluorescence appeared throughout the spheroid (Figure 5A). To determine viability of these spheroids following PFI-3 treatment, we used the MTS assay. In MTSs with ARID1A, a steady but small decline in viability was observed with increasing concentration of the drug with significant loss in viability observed only at high concentrations (above 200 μM PFI-3) (Figure 5B). ARID1A^-/-^ MTSs on the other hand showed increased sensitivity to the drug with significant loss in viability at concentrations above 100 μM (Figure 5B). IC _50_value of PFI-3 was calculated using Hills equation. The IC _50_value of PFI-3 for HCT116 MTS was calculated to be 300μM while those of MTS lacking ARID1A was 99μM suggesting that loss of ARID1A made MTS 3 times more sensitive to PBRM1 inhibitors. Our observations indicate that PBRM1 does play important roles in spheroids lacking ARID1A and that loss of PBRM1 represents a new synthetic vulnerability within cancers arising from ARID1A loss.

**Figure 5:**
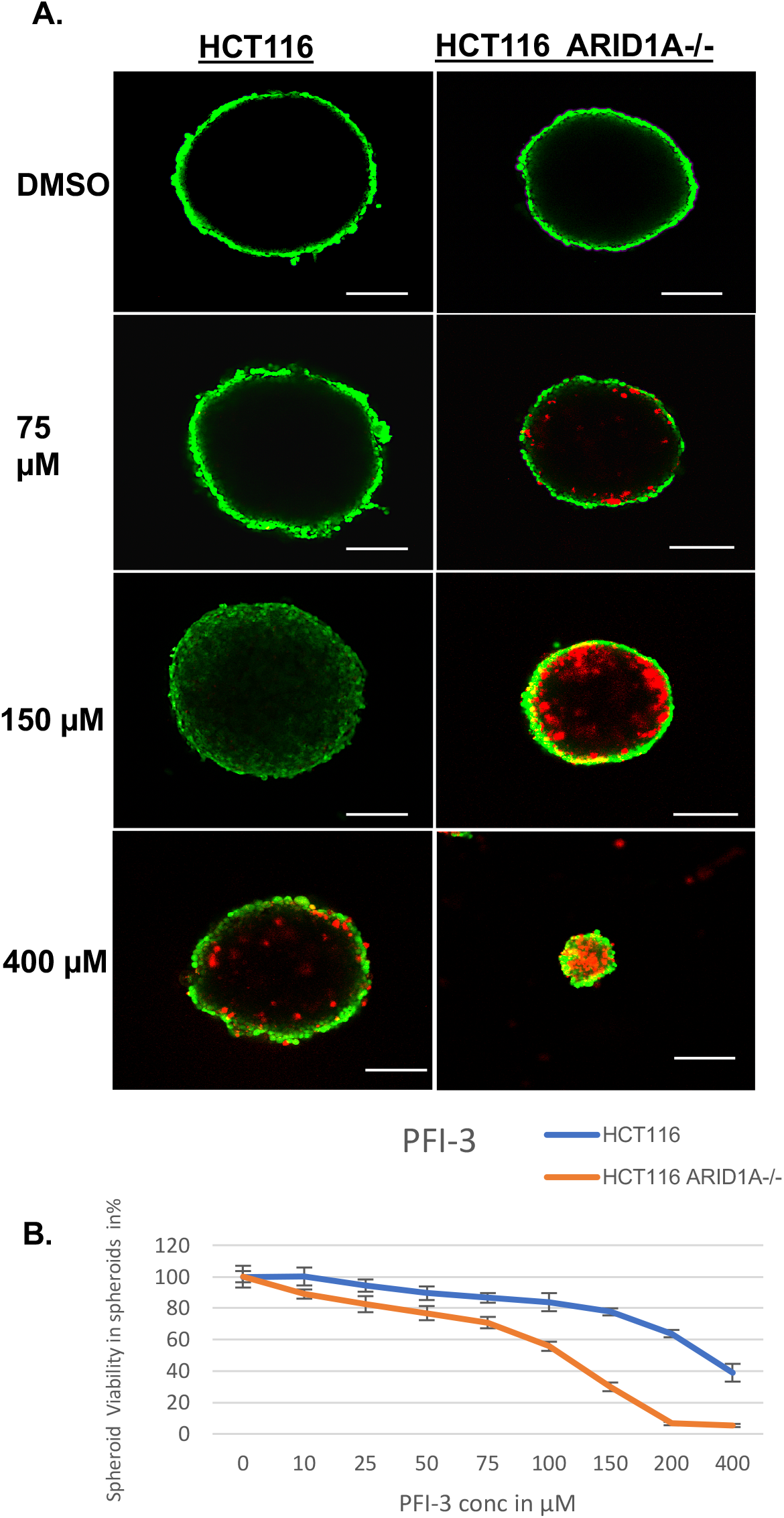
HCT116 ARIDA-/- MTS display increased sensitivity to PFI-3. **A)** Cells were seeded and allowed to form MTS for 72 hours. MTS were treated with PFI-3 for 72 hours and viability using a live/dead staining with ethidium homodimer-1 and Calcein AM. Viable cells appear as green, while dead cells appear as red. Images represent a section of Z stack taken using confocal microscope. Scale bars represent 100 μm. **B)** Quantification of viability of PFI-3 treated MTSs using Promega 3D Cell Titre GLO assay. Error bars represent SD of three independent experiments.

## Discussion

The increase in mutational frequency of SWI/SNF subunits in human diseases and one of the most frequently mutated tumor suppressors in cancers, has led to identification of vulnerabilities of the chromatin regulator. Vulnerabilities have focused mainly on mutually exclusive subunits due to residual SWI/SNF complexes present. Most of these studies have been demonstrated in 2D culture and reported ARID1B as a synthetic lethality in ARID1A-deficient cells.

Unlike 2D models, a 3D spheroid is more representative of a tumor model, but the vulnerabilities of SWI/SNF in 3D models have yet to be explored. Here, we uncovered that in the 3D model, loss of unique subunits, ARID1B, PBRM1, BRD7, and BRD9 did not have a significant impact on spheroid formation or viability in HCT116. However, these same knockdowns inHCT116 ARID1A-/-, loss of ARID1B and PBRM1 showed significant defects in formation (Figure 2A), survival (Figure 2B) and proliferation (Figure 2C). Interestingly, loss of BRD9, a subunit of ncBAF, remained proliferative and viable with loss of ARID1A, suggesting that ARID1B-BAF and PBAF can replace ARID1A-BAF for tumor survival. Our observations of PBRM1 represents a previously undescribed vulnerability within SWI/SNF on loss of ARID1A in spheroids and not in 2D cell culture. Our result reveals that in addition to ARID1B, PBRM1 could be targeted in tumors lacking ARID1A.

In addition to identifying ARID1B and PBRM1 as a vulnerability in 3D spheroids lacking ARID1A, we also characterized the composition of SWI/SNF in these spheroids. WT MTSs contains three complexes, ARID1A-BAF, PBAF, and ncBAF, with reduced levels of ARID1B-BAF (Figure 3A). However, in MTSs lacking ARID1A, there is a shift in the balance of complexes with increased predominance of ARID1B-BAF and PBAF, with ~ 4-fold increase in PBRM1 (Figure 3A). ncBAF though present has lower levels of BRD9 (Figure 3A). We confirmed this with glycerol sedimentation fractionation, showing thatHCT116 ARID1A-/- spheroids rely on ARID1B-BAF and PBAF (Figure 3B). Our studies also show the importance of ARID1B and PBRM1 in spheroids lacking ARID1A in induction of hypoxic genes. We show that ARID1B and PBRM1 are required for SWI/SNF-dependent hypoxic genes PAI1 and EPO but SWI/SNF-independent hypoxic gene, GLUT1, continues to not depend on SWI/SNF (Figure 4).

As there are very few inhibitors that can target ARID1B, we decided to target PBRM1 in the 3D model. We used PFI-3 that selectively binds to the fifth bromodomain of PBRM1.^42,43^ Treatment with PFI-3 increased death within ARID1A^-/-^ MTSs but not significantly in wild type MTSs at low concentrations of the drug (Figure 5A). We also tested how the viability of the spheroids would be affected with PFI-3. In both spheroid models, PFI-3 slowly decreased viability, but the effects were noticeable for HCT116 spheroids at a concentration above 200 μM and for HCT116 ARID1A-/-, significant affects were visible at around 50-75μM (Figure 5B). Hills equation revealed that MTS lacking ARID1A are 3 times more sensitive to PBRM1 inhibitors. These results confirm PBRM1 as a vulnerability in spheroids lacking ARID1A.

We have shown here for the first time the heterogeneity of SWI/SNF in a 3D model MTS system. Our work confirms ARID1B as a vulnerability in ARID1A-deficient cells, but we reveal PBRM1 as an added vulnerability on loss of ARID1A. Our studies show that both ARID1B and PBRM1 are needed to replace the functions of ARID1A in ARID1A deficient tumors and allows opportunities to explore the development of new therapeutic strategies to target cancers arising from loss of ARID1A.

## Materials and Methods

### Cell Culture

HCT116 and HCT116 ARID1A^-/-^ line (generated by a knock-in of a premature stop codon (Q456*)) were purchased from Horizon Discovery and maintained in McCoy’s medium supplemented with 10% FBS (Gibco) and 1% penicillin/streptomycin (Gibco). All cells were tested for mycoplasma (Sigma) before the start of experiment. For normoxic cell culture, cells were cultured in a 5% CO_2_ humidified incubator at 37°C.

### Nuclear Extract Preparation

Extracts were prepared using lysis buffer consisting of 20 mM HEPES, pH 7.9, 1.5 mM MgCl_2_, 420 mM KCl, 0.2 mM EDTA, 25% glycerol, 0.1 mM DTT, complete protease inhibitors, and 1 mM PMSF. The buffer was supplemented with benzonase and heparin to aid in extraction of chromatin bound proteins. Protein concentration was determined by BCA (Thermo Fisher). For western blotting, equal amounts of nuclear extracts were loaded onto SDS PAGE gel and subjected to immunoblotting with protein specific antibody. The following primary antibodies were used: anti-ARID1A (Sigma Aldrich: HPA005456), anti-ARID1B (Bethyl Laboratories: A301-046A), anti-BRD9 (Bethyl Laboratories: A303-781A), anti-BAF155 (Santa Cruz: sc-9746), anti-anti-BRG1 (Abcam: ab110649), anti-GLTSCR1 (Gen Script), anti-BAF170 (Bethyl Laboratories: A301-039A), anti-BAF60a (Santa Cruz: sc-514400), anti-BAF57 (Bethyl Laboratories: A300-810A), anti-INI1 (Bethyl Laboratories: A301-087A), anti-PBRM1 (Sigma Aldrich: HPA015629), anti-BRD7 (Proteintech: 51009-2-AP), and anti-TBP (Proteintech: 22006-1-AP).

### siRNA Transfections

500,000 cells were seeded into 6-well plates and incubated overnight. A double transfection was performed using siRNA cocktail (ON-TARGET Plus Pool, Horizon Discovery). 10 μl of 5 μM siRNA was diluted into 250 μl of Opti-MEM (Gibco) and 10 μl of Lipofectamine 2000 (Invitrogen) was diluted into 250 μl of Opti-MEM. The solutions were incubated at room temperature for 5 minutes and then mixed and incubated at room temperature for 20 minutes. The mixture was then added dropwise in each well and incubated at 37°C. Another round of transfection was performed 24 hours later. The cells were harvested after 48 hours. Spheroid experiments and cell viability assays were performed 24 hours after second transfection.

### Immunoprecipitation

Approximately 2.5 μg of BRG1 antibody (Abcam) and 20 μl of Protein A Dynabeads (Invitrogen) were crosslinked at 4°C, rotating end over end for 4 hours. Antibody-bound beads were washed 4X with ice cold PBS and once with 1 ml 0.1 M sodium borate (pH 9.0). Antibody-bound beads were resuspended with sodium borate with the addition of 5.2 mg of dimethylpyrimilidate (DMP) and allowed to rotate end over end at room temperature for 30 minutes. Antibody-bound beads were washed once with 0.2 M ethanolamine (pH 8.0) and then once more and allowed to rotate end over end at 4°C overnight, prior to nuclear extraction. Nuclear extract preparation was used for IP. Approximately 1 mg of protein extract was added to Buffer containing 20 mM HEPES, pH 7.9,1.5 mM MgCl2, 300 mM KCl, 0.2 mM EDTA, 10% glycerol, 0.1% NP-40 with complete protease inhibitors, 0.1 mM DTT and 1 mM PMSF and incubated on ice for 15 minutes and spun at 14,000 rpm for 20 minutes at 4°C. Supernatant was removed and added to antibody-bound beads and incubated at 4°C, rotating end over end, for 4 hours. Nuclear extracts were eluted in 0.1 M glycine (pH 2.5) for 2 minutes on ice with constant flicking of tube. Tris-HCl pH 8.8 was added to neutralize the pH.

To verify quality of samples, silver staining and western blotting was performed. The SDS-PAGE was fixed in Buffer 1 (50% ethanol, 10% acetic acid and 40% deionized water) overnight and washed 3X for 15 minutes each thereafter. Gels were sensitized in Buffer 2 (0.02 g of Na2S2O3 in 100 ml of deionized water) for 25 minutes, washed 3X with deionized water and allowed to develop in Buffer 4 (6 g of Na2CO3, 2 ml of Na2S2O3, 50 μl of formaldehyde in 100 ml of deionized water) until a desired signal and stopped by stop solution (10% acetic acid in 100 ml of deionized water) for 20 minutes.

### Evaluation of cell viability in monolayers

8,000 cells of each treatment were diluted in 200 μl of media and seeded in a 96-well plate and allowed to incubate overnight. Following overnight incubation, media was changed in the wells. To assay cell viability, MTS assay was performed by adding 20 μl of reagent (Promega CellTiter 96 Aqueous One Solution) to each well and incubated at 37°C for 2 hours. Absorbance was measured at 490 nM (BioTek).

### Generation of spheroids

Molds from Microtissues Inc. were used to make scaffolds. 2% agarose solution was poured into the molds and allowed to solidify. The scaffolds were then isolated in 6 well plates and sterilized under UV for 2 hours. The scaffolds were seeded with 150,000 cells diluted in 100 μl of media, incubated at 37°C and monitored for spheroid formation.

### Cell labeling and imaging of MTS

150,000 cells of each treatment were seeded and incubated for 96 hours. The MTS were stained with a mixture of 2 μM calcein AM and 4 μM ethidium homodimer-1 in PBS (Thermo Fisher Scientific) to stain for live and dead cells, respectively. The MTS were then fixed with 4% paraformaldehyde and then 2% sucrose. The slide was incubated for an hour. Brightfield and fluorescent images were captured using confocal microscopy (Nikon Confocal).

### 3D cell viability assay

150,000 cells were seeded in the molds and incubated in the incubator. At the end of Day 3 and Day 7, MTS formed were harvested and transferred to a 96-well plate. 100 μl of media and 100 μl of reagent (3D CellTiter Glo) were added to each well, according to the manufacturer’s instructions. After 30 minutes of incubation at room temperature, the total luminescence values were measure using a multiplate reader (Syngene).

### Glycerol Gradient Centrifugation

1 mg of nuclear extract supplemented with protease inhibitors was carefully overlaid onto a 10 ml 10-30% glycerol gradient prepared in a polyallomer centrifuge tube (Beckman Coulter). Tubes were placed in a SW-41 swing bucket rotor and centrifuged at 4°C for 16 hours at 40,000 rpm. 500 μl of 100 mM Tris-HCl, pH 8.5 and 250 μl TCA were added to fractions, inverted, and placed into 4°C, overnight. Factions were spun at 14,000 rpm for 30 minutes at 4°C, supernatant removed and washed twice with 500 μl of cold acetone, inverting gently. Supernatant was removed after washes and allowed to air dry in the hood. Fractions were then subjected to western blotting analyses.

### Mass Spectrometry

For mass spectrometry, complexes were purified by Immunoprecipitation and precipitated by TCA method. TCA precipitated samples were sent to IDeA National Resource for Quantitative Proteomics, Arkansas.

### Quantitative RT-PCR analysis

RNA from siRNA treated MTS were obtained using Trizol (Invitrogen). cDNA was synthesized using random hexamer by SuperScript IV First-Strand cDNA Synthesis System (Thermo Fisher). The primers for quantitative RT-PCR analysis are listed. (Supplementary Figure 5). Quantitative real-time PCR was performed with Power SYBR Green PCR Master Mix (Applied Biosystems). The relative amount of target gene mRNA was normalized to 18S mRNA by the ΔΔCT method.

## Supporting information

all supplemental figures

## Acknowledgements

We would like to thank all members of the Dutta and Menon labs for their help and comments. This work is supported by funding to AD from the Rhode Island Foundation Medical Research Grant, <u>NIH/NIGMS P20GM103430/Rhode Island IDeA Network for Biomedical Research Excellence (RI-INBRE)</u> and the University of Rhode Island.

