## Supplementary material for "PBRM1 is a vulnerability in ARID1A deficient multicellular tumor spheroids": all supplemental figures

### Supplementary data

**A.**

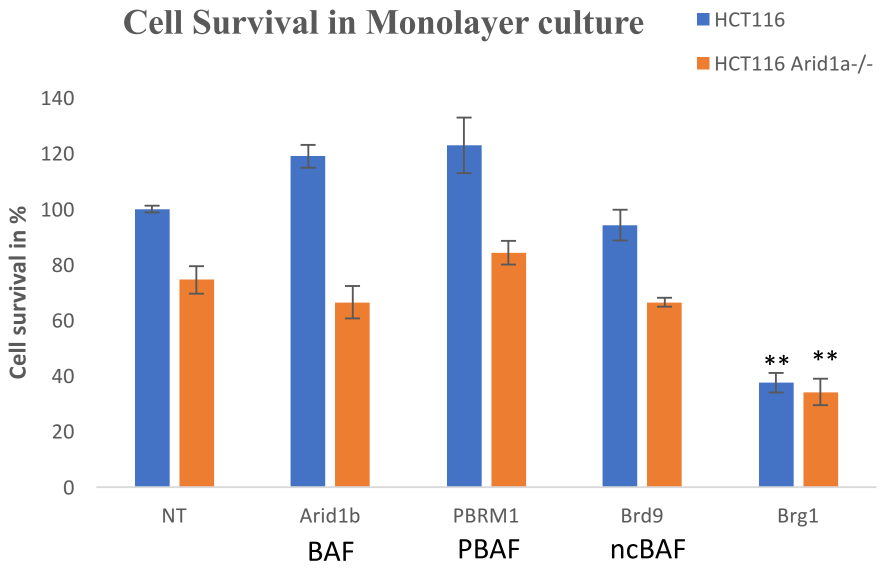


**B.**

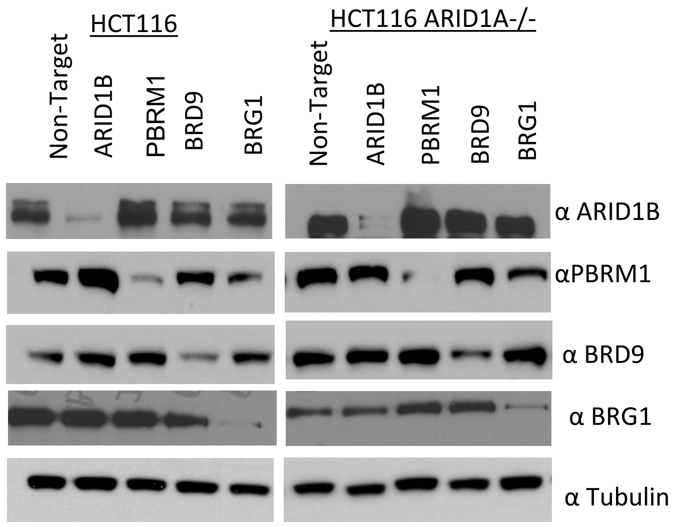


**Sup Figure 1: Cell survival of HCT116 and HCT116 Arid1a-/- grown in monolayer. A)** Cells were treated with siRNA and seeded onto 96 well plates and viability of cells was quantified as a measure of ATP activity Survival of HCT116 non target cells were used as reference for normalization. **B)** Western blot images of HCT116 and HCT116 Arid1a-/- cells confirming knockdown of proteins. Total protein was isolated from cells grown in monolayer after treatment with siRNA. Loss of proteins was confirmed by probing for protein with antibody specific to different subunits by western blotting. α Tubulin levels was used as loading control for the experiment. ***<0.05

A.

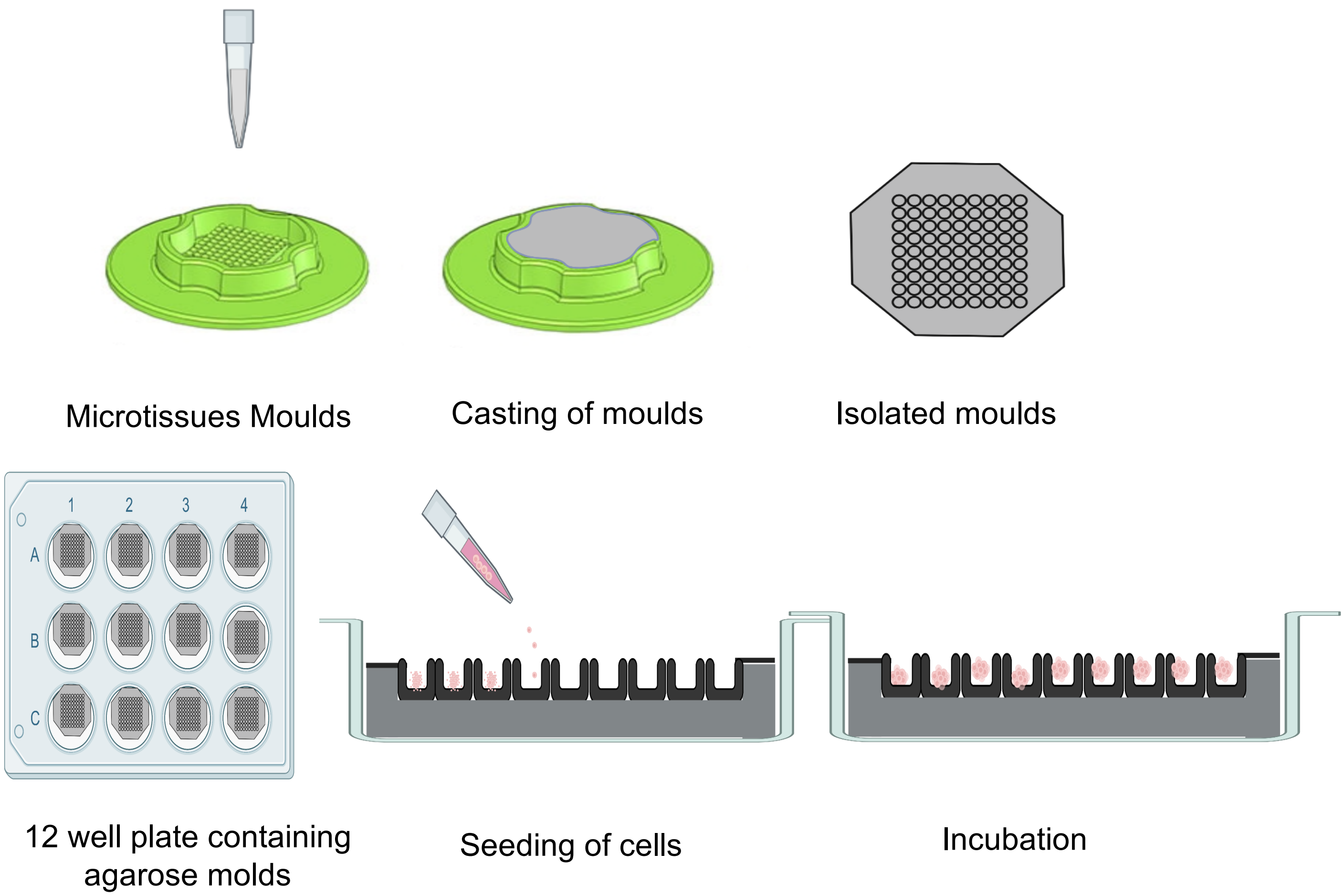


B.

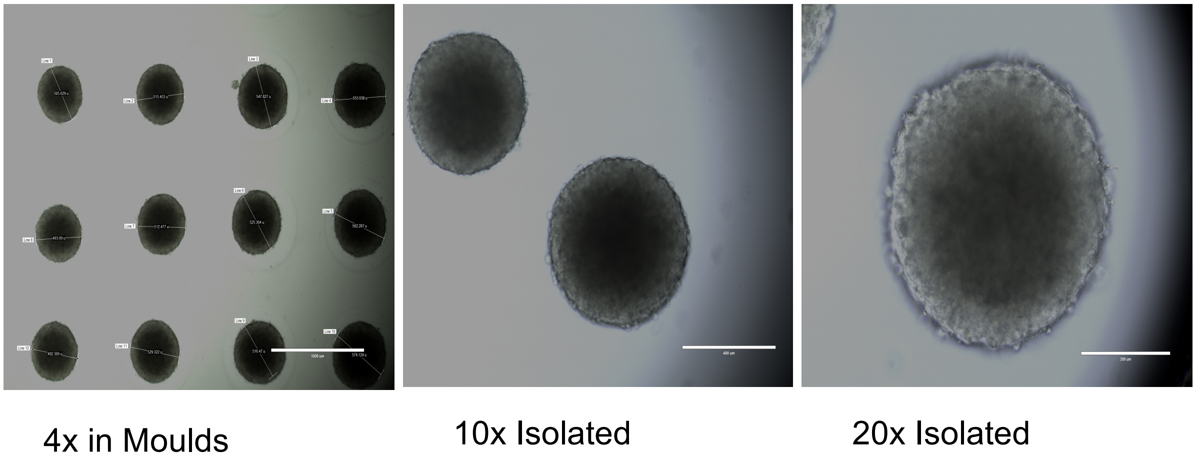


**Sup Figure 2**: **Generation of tumorspheres. A)** Microtissues molds were used as scaffolds to cast agarose molds. Agarose was poured into these molds and allowed to set. Cells were then seeded and incubated for 72 hours. **B)** Representative brightfield images of Tumorspheres at 4X, 10 X and 20X magnification. Scale bars represent 400 µm.

A.

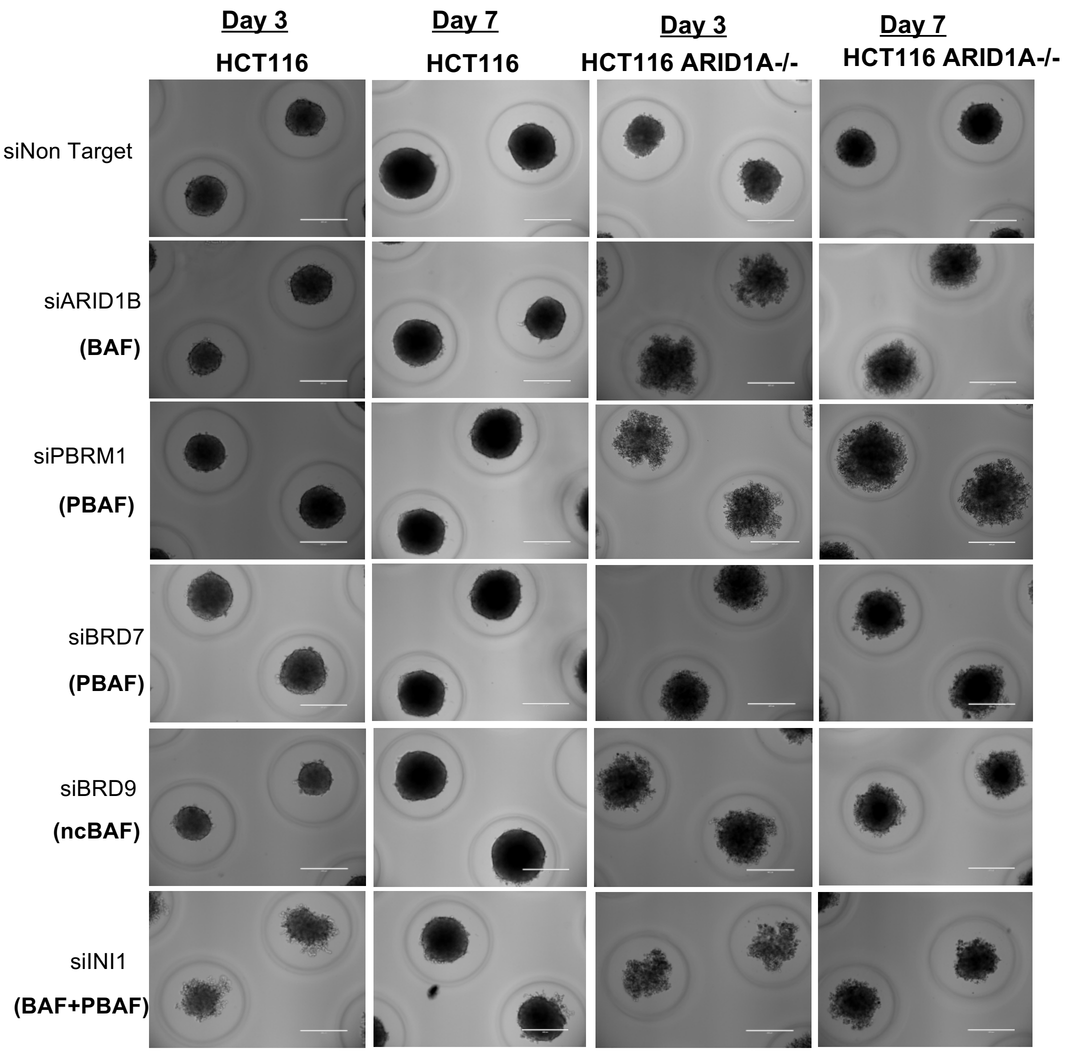


B.

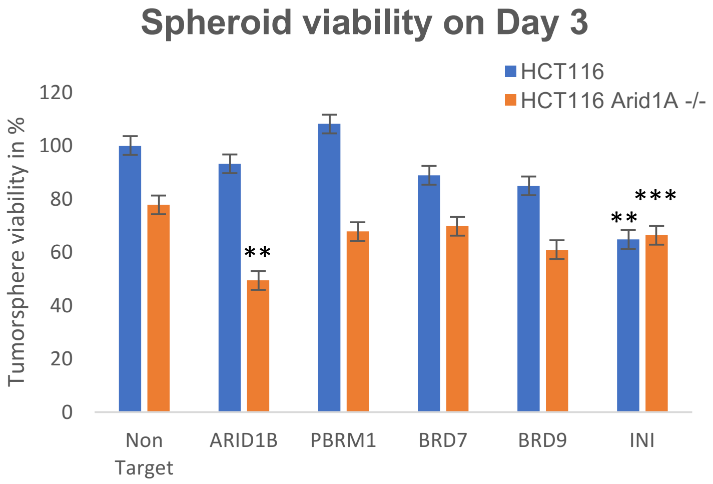


**Sup Figure 3 : Quantification of viability in Multicellular Tumorspheres lacking ARID1A-/- upon loss of SWI/SNF subunits.** HCT116 and HCT116 ARID1A-/- cells were treated with siRNA against subunits of SWI/SNF and allowed to form tumorspheres. The tumorspheres were imaged regularly and quantified for viability over 7 days.. Scale bars represent 100 µm. **A)** Brightfield images of tumorspheres treated with siRNA taken on day 3 and day 7 using EVOS microscope. Scale bars represent 400 µm. **B)** Graphs showing quantification of viability of HCT116 and HCT116 ARID1A-/- MTS lacking SWI/SNF subunits on day 3 with Promega 3D Cell Titre GLO. Error bars represent SD of three independent experiments. ***<0.05

A.

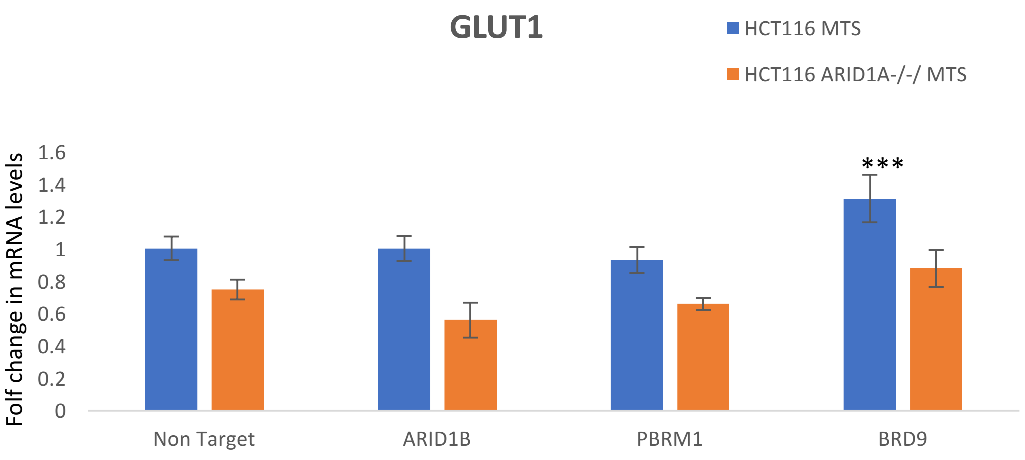


B.

| Primer | Sequence (5’-3’) |
|---|---|
| 18S Forward | GAATTCCCAGTAAGTGCGGG |
| 18S Reverse | GGGCAGGGACTTAATCAACG |
| EPO Forward | TTCGCAGCCTCACCACTCT |
| EPO Reverse | GAGATGGCTTCCTTCTGGGC |
| PAI1 Forward | AAGGCAACATGACCAGGCT |
| PAI1 Reverse | GGGAGAACTTGGGCAGAACC |
| ANGPTL4 Forward | TCCGTACCCTTCTCCACTTG |
| ANGPTL4 Reverse | AGTACTGGCCGTTGAGGTTG |
| BNIP3LForward | CTGTACAGTCTTCCCAAGGTG |
| BNIP3L Reverse | GCTACATGAGAAAATGACCAG |
| GLUT1 Forward | ATTGGCTCCGGTATCGTCAAC |
| GLUT1 Reverse | GCTCAGATAGGACATCCAGGGTA |

**Sup Figure 4 :Multicellular tumorsphere qRT-PCR. A)** qRT-PCR analysis of the mRNA levels of SWI/SNF independent hypoxia marker- GLUT1 in HCT116 and HCT116 ARID1A-/- MTS. Cells were treated with siRNA against specific subunits and allowed to form MTS for 72 hours. RNA was isolated from the spheroids and mRNA quantified to calculate fold changes in induction. Error bars represent SD of three independent experiments. **B)** List of primers used in the study for qRT-PCR. ***<0.05

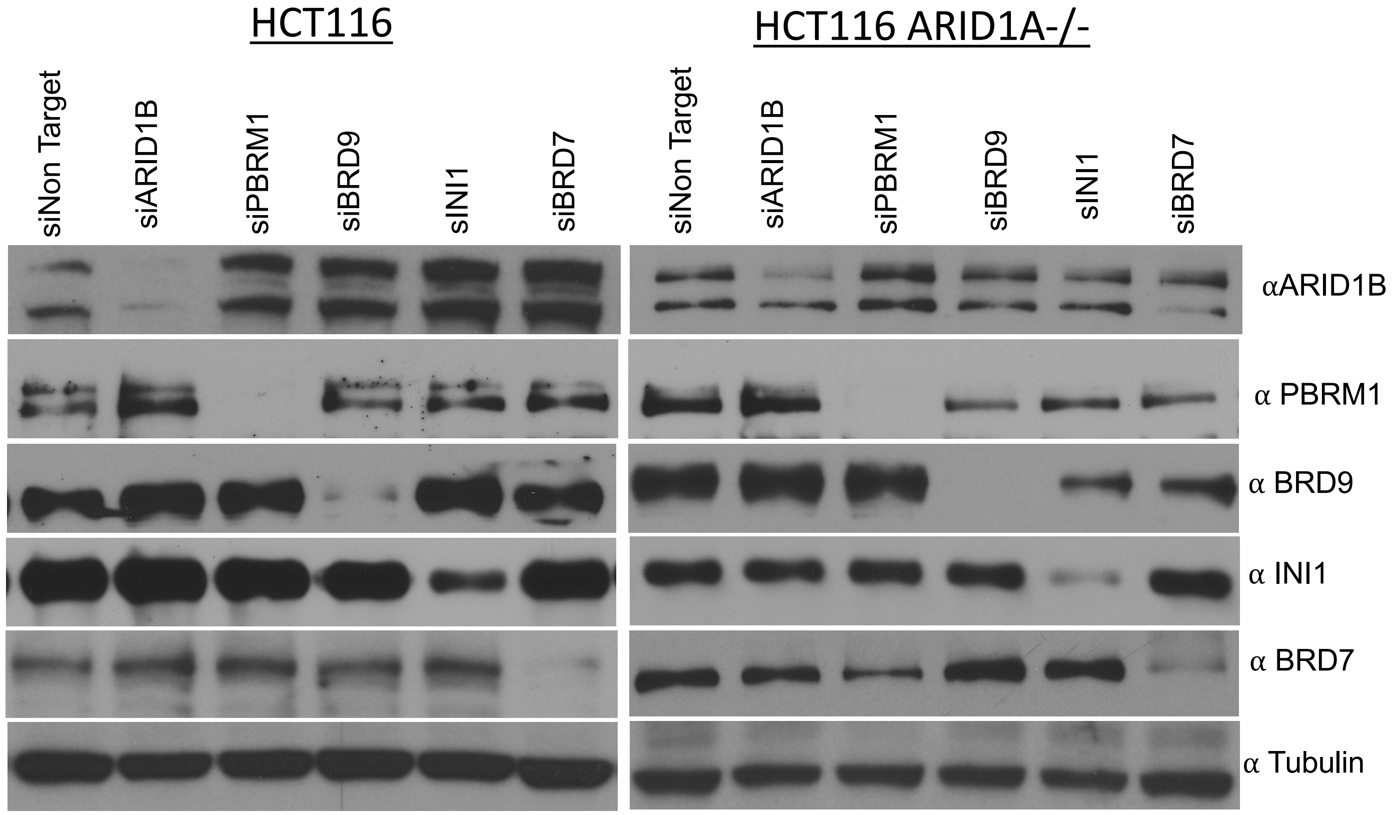


**Sup Figure 5: Western blot images of HCT116 and HCT116 ARID1A-/- cells confirming knockdown of SWI/SNF proteins in cells seeded for Calcein /Ethidium Homodimer 1 assay.** Cells were treated with siRNA. 24 hours post treatment,150000 cells/well were seeded to form tumorspheres while remaining cells were cultured as monolayer culture and harvested after 4 days for checking protein levels. Loss of proteins was confirmed by probing for protein with antibody specific to different subunits by western blotting. α Tubulin levels was used as loading control for the experiment.
